# The influence of positive emotion and negative emotion on false memory based on EEG signal analysis

**DOI:** 10.1101/2021.01.12.426168

**Authors:** Ying Li, Zhaobing Ni, Renjie He, Junyu Zhang, Zhimou Zhang, Shuo Yang, Ning Yin

**Author notes:** Corresponding author at: School of Electrical Engineering, Hebei University of Technology, Tianjin, CA 300130, China. *E-mail address:* (Y. Li).

## Abstract

Analyzing the influence of emotion on false memory through electroencephalogram is helpful to further explore the cognition function of brain. In this study, we improved the Deese-Roediger-McDermott paradigm experiment to study the false memory. The memory materials are combined with mixed emotions, which are closer to real life. Twenty-eight participants were randomly divided into positive group and negative group. We used music to induce the participants in the positive group and the negative group to generate corresponding emotions. Finally, we analyzed the difference between the positive group and the negative group from the behavior data, source location and cortex functional network of event related potential. The results of behavioral data analysis show that the false memory rate of positive group (85%) is significantly higher than that of negative group (72%). The results of source localization show that the brain of the positive group is more active than that of the negative group, and the difference of brain activation location between the two groups is mainly manifested in the BA24 and BA45 brain regions. The results of cortex functional network show that the node degree, clustering coefficient, global efficiency and small-world property of the positive group are significantly higher than those of the negative group. It can be found from the three aspects that participants’ emotional state and brain’s understanding of semantic are the main reasons for the difference in the incidence of false memory between the two groups.

## 1. Introduction

The process of memorizing, recognizing or maintaining the events experienced by the brain is a process of memory, but this process is not always correct. It may produce errors spontaneously or due to external interference. When people recall or recognize some events that they have not seen, they mistakenly think that they have seen the events, and then false memory occurs. False memories, like correct memories, appear around us almost every day [1, 2].

In recent years, along with the researches on the influence of emotion on correct memory, many scholars have begun to study the relationship between emotion and false memory. At present, the influence of emotion on memory is often studied from two aspects: memory materials with emotion and emotional state of participants. In the past researches, some researchers asked participants to memorize emotional vocabulary in the neutral emotional state. They found that participants had more false memories of words with negative emotions than words with positive emotions [3, 4]. Some researchers used neutral vocabulary to study the influence of participants’ emotional state on false memory, and found that participants in positive emotional state produced more false memories than those in negative emotional state [5, 6]. Some researchers combined emotional vocabulary with participants’ emotional state, and found that there was also emotional consistency in false memory [7]. For example, participants in positive emotional state would produce more false memories of words with positive emotions. After sorting out the previous studies on false memory, it is found that most of them are based on behavioral data or psychological analysis, and only a few researchers have studied the electroencephalogram (EEG) signals of false memory [8, 9].

As a typical bioelectrical signal, EEG signal contains a lot of pathological or physiological information. The study of EEG signal can not only diagnose some brain diseases clinically, but also help us to explore the working mechanism of brain essentially under non-invasive conditions. The EEG signals collected on the scalp have high time resolution, but the interference caused by scalp, skull and cerebrospinal fluid is inevitable. Therefore, estimating the position and intensity of neural activity sources in the brain based on the EEG signals recorded on the scalp can help us understand the working mechanism of the brain more accurately [10]. The standardized low resolution brain electromagnetic tomography algorithm (sLORETA) is a linear source localization algorithm proposed by Pascal-Marqui [11]. When sLORETA is used to locate the source of EEG signal, the spatial resolution will decrease with the increase of depth, but the linear characteristic of the algorithm will make the error of the source position be zero in the ideal case [12–14]. It uses finite inversion to estimate the probability sources of EEG signals in the standard brain atlas space. At present, it has been widely used in the detection of Alzheimer’s disease, epilepsy, depression and other brain diseases [15–17].

There are many complex networks which are independent and influence each other in our life. By combining complex networks with brain neuroscience, we can study the cognitive activities of the brain in time domain or frequency domain through some characteristics of brain networks. In recent years, the construction and analysis of brain function network has been widely used in the research of mental disease, neurological disease, behavioral disorder and other brain diseases [18–20]. However, the spatial resolution of brain network directly established by scalp EEG signal is low and has the effect of cranial cavity volume effect. Therefore, some researchers reconstruct the source from the scalp EEG signals firstly, map the localization to the cortex, and then construct the functional network of cortex. This method improves the spatial-temporal resolution and makes the constructed brain network more reliable [21, 22].

People’s emotions will change when they are affected by external factors. In the process of learning and memory, the emotion of the memory material that people face is also changeable. In order to get closer to real life, we designed Deese-Roediger-McDermott (DRM) paradigm experiment using memory materials with mixed emotions. DRM paradigm experiment can well induce participants’ false memory and record it [23]. By comparing the source localization results of event-related potential (ERP) of false memory and the characteristics of functional network of cortex, we analyzed the influence of participants’ emotional state on false memory in DRM paradigm experiment with mixed emotional memory materials. The whole research process is shown in Fig. 1. This study provides a theoretical basis for how to avoid the occurrence of false memory in life and to study the working mechanism of brain memory.

**Fig. 1.**
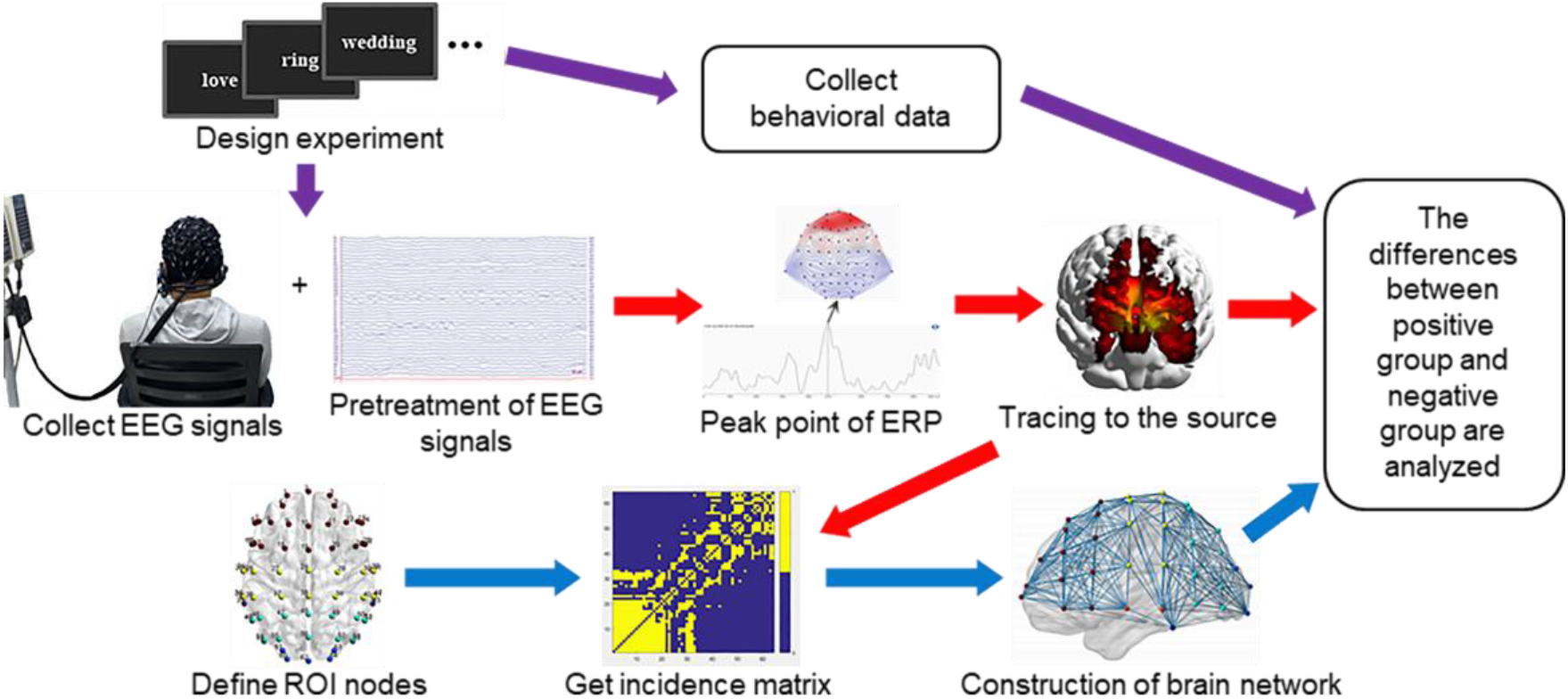
Flow chart of research sequence. The purple arrow indicates the behavior data difference analysis process. The red arrow indicates the source location difference analysis process. The blue arrow indicates the brain network difference analysis process.

## 2. Materials and methods

### 2.1. Experimental

In this study, E-Prime 3.0 software was used to design the experimental interface and record the behavior data of participants in the background. Neuroscan equipment was used to collect 64 channel EEG signals. The distribution of electrode channels conformed to the international 10-20 electrode lead positioning standard. The sampling frequency was 1 000 Hz. The impedance between the electrode and the scalp was kept below 10 kΩ. In the experiment, music that elicited the participants’ emotions was played through headphones.

Twenty-eight 22 to 25 years old graduate students (fourteen females) participated in the experiment. The participants were chosen as right-handed, native Chinese speakers, with no history of mood disorders and depression. Participants were randomly assigned to positive or negative mood conditions (seven females and seven males in each condition). Mood was induced using music. Participants in the positive mood-induction group listened to 3 minutes of Bach’s *Brandenburg Concerto No. 3* and participants in the negative mood-induction group listened to 3 minutes of Prokofiev’s *Alexander Nevsky: Russia Under the Mongolian Yoke*. The experiment was approved by the Biomedical Ethics Committee of Hebei University of Technology (approval number HEBUThMEC2020006). No ethical issues were involved in the experiment, and the study conformed to the Declaration of Helsinki. All participants signed the informed consent before the experiment.

An improved DRM paradigm was used to induce participants’ false memories. In our experimental materials, different from other researchers, 12 emotional word lists (four positive, four neutral, four negative) were mixed and used. Each word list contained 12 learning words (all of them appeared in the learning phase, some of them appeared in the test phase) and 1 keyword was used to induce false memory (it only appeared in the test phase to induce false memories). In the learning phase, the lists were mixed and presented to the participants. In the test phase, the error rate of these keywords was the false memory rate of participants. Before the learning stage and the test stage, the participants’ emotions were fully induced. During the learning phase, each word was displayed on the computer monitor for 1s, and the word needs to be remembered by the participants. During the test phase, each word was displayed on the computer monitor for 2s. If the participants think they had seen the word in the learning phase, they were asked to press the “F” key on the keyboard with their left index finger, and otherwise press the “J” key on the keyboard with the right index finger.

### 2.2. Source location

After preprocessing the collected EEG signals, we superposed and averaged the EEG signals of false memory, and obtained the ERP of participants’ false memory. We used standard low resolution brain electromagnetic tomography algorithm (sLORETA) to locate the source of ERP [11]. A real head boundary element model was used. The distance between voxels in the head model was 2.5 mm, and the source localization results were constrained on the cerebral cortex. The distribution of current density on the cerebral cortex can be obtained. We found a significant difference between the positive group and the negative group. The peak values of current density in positive group and in negative group appeared at 474 ms and 820 ms, respectively. We took the peak point as the zero point and selected ERP from −100 ms to 100 ms for source localization. In order to study the difference of source location more accurately, we used the statistical nonparametric mapping (SnPM) method [24] to analyze the difference of source location between the positive group and the negative group, and the difference results are displayed in Brodmann (BA) brain area.

### 2.3. Cortex functional network

To construct a cortex functional network, the nodes and edges must be determined first. We defined 64 regions of interest (ROI) on the cerebral cortex and used the central points of these regions as nodes. These nodes correspond to the electrode points on the electrode cap and spread all over the cerebral cortex.

The degree of association between two nodes in the network is the connection edge. We used sLORETA to locate the source of the EEG signals. Finally, we get 31061 voxel points per millisecond in the cerebral cortex, and each voxel point has a corresponding current density value. We redistributed the 31061 current density values to 64 ROI nodes according to the principle of proximity, and obtained the current density values of 64 ROI nodes per millisecond. The correlation matrix of brain network can be constructed by using the current density value of ROI node. We used the Pearson cross correlation algorithm to calculate the degree of association between every two ROI nodes, and got the 64×64 correlation matrix.

The appropriate threshold was selected to binarize the correlation matrix, and then the brain network was constructed by using the binary matrix. When selecting the threshold, we should not only ensure the integrity of the network, but also avoid the network being too dense, and at the same time ensure the small-world property of the network. When the average node degree 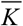 of the network is larger than the natural logarithm of the node number *N*, that is, 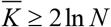, it can meet the above requirements. After calculation, we choose the threshold value as *T* = 0.85.

### 2.4. Brain network analysis method

We analyzed the topological structure of the cortical functional network from the node degree, clustering coefficient, global efficiency and small-world property.

#### 2.4.1. Node degree

In the network, the node degree represents the total number of nodes connected with node *i*, which is a basic measurement index in complex networks. The greater the degree of nodes, the more connected edges and the more important it is in the network [25]. The calculation formula of node degree is as follows:

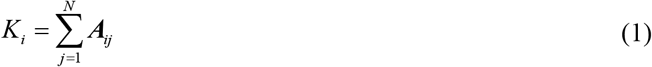

where *N* is the total number of network nodes, *A_ij_* is the element in the binary matrix.

#### 2.4.2. Clustering coefficient

In the network, the clustering coefficient represents the ratio of the actual number of adjacent nodes edges of node *i* to the number of all possible adjacent nodes edges. It is used to measure the clustering characteristics and compactness of the network, and can reflect the connectivity of brain functional network [26]. The calculation formula of clustering coefficient is as follows:

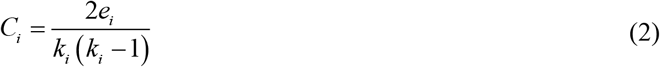

where *e_i_* is the actual number of adjacent nodes of node *i*, *k_i_* (*k_i_* − 1)/2 is the number of possible edges.

The clustering coefficient of the whole network is represented by the average value of clustering coefficient of all nodes in the network. The average clustering coefficient *C* ranges from 0 to 1, and the larger its value, the closer the connection between brain network nodes. The calculation formula of average clustering coefficient is as follows:

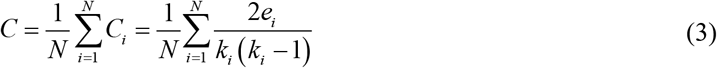

#### 2.4.3. Global efficiency

In the network, the global efficiency represents the average of the reciprocal of the shortest path length. The shortest path length and global efficiency can be used to measure the global transmission ability of the network. The shorter the shortest path length, the higher the global efficiency of the network, and the faster the rate of information transmission between nodes in brain network [27]. The calculation formula of global efficiency is as follows:

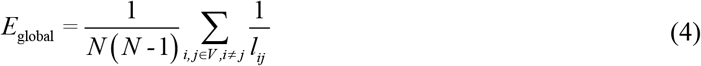

where *V* is the set of all nodes, *N* is the total number of network nodes, *l_ij_* is the path length between node *i* and *j*.

#### 2.4.4. Small-world property

Small-world network represents a network with high clustering coefficient and shortest path length. Small-world property *σ* of the network can be quantitatively analyzed [28], and its description is as follows:

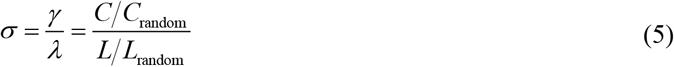

where *C* is the average clustering coefficient of the network, *C*_random_ is the clustering coefficient of a random network with the same number of nodes and edges as the network, *L* is the average path length of the network, the network. *L*_random_ is the path length of a random network with the same number of nodes and edges as the network.

When *σ* > 1, it means that the network has small-world property, and the larger the value of *σ*, the stronger the small-world property of the network.

## 3. Results

### 3.1. Behavioral Data

We counted the number of keystrokes participants made. The results showed that the average false memory rate was 85% in the positive group and 72% in the negative group. We used independent sample t-test in SPSS 20.0 statistical software to test the false memory rate of the two groups (*p*=3×10^−3^<0.05). It showed that the false memory rate of positive group was significantly higher than that of negative group.

### 3.2. Source location

The source localization results are shown in Fig. 2. By analyzing the source localization results, we find that the positive group is mainly located in the frontal and temporal lobes (see Fig. 2(a)). The negative group is mainly located in the prefrontal and right temporal lobes (see Fig. 2(b)). The activation intensity of the positive group is higher than that of the negative group. Combined with the behavioral data, we find that the positive group has higher activation intensity and more false memories. The negative group has lower activation intensity and less false memories. The results show that the more active the brain is, and the more false memories are produced when identifying key words.

**Fig. 2.**
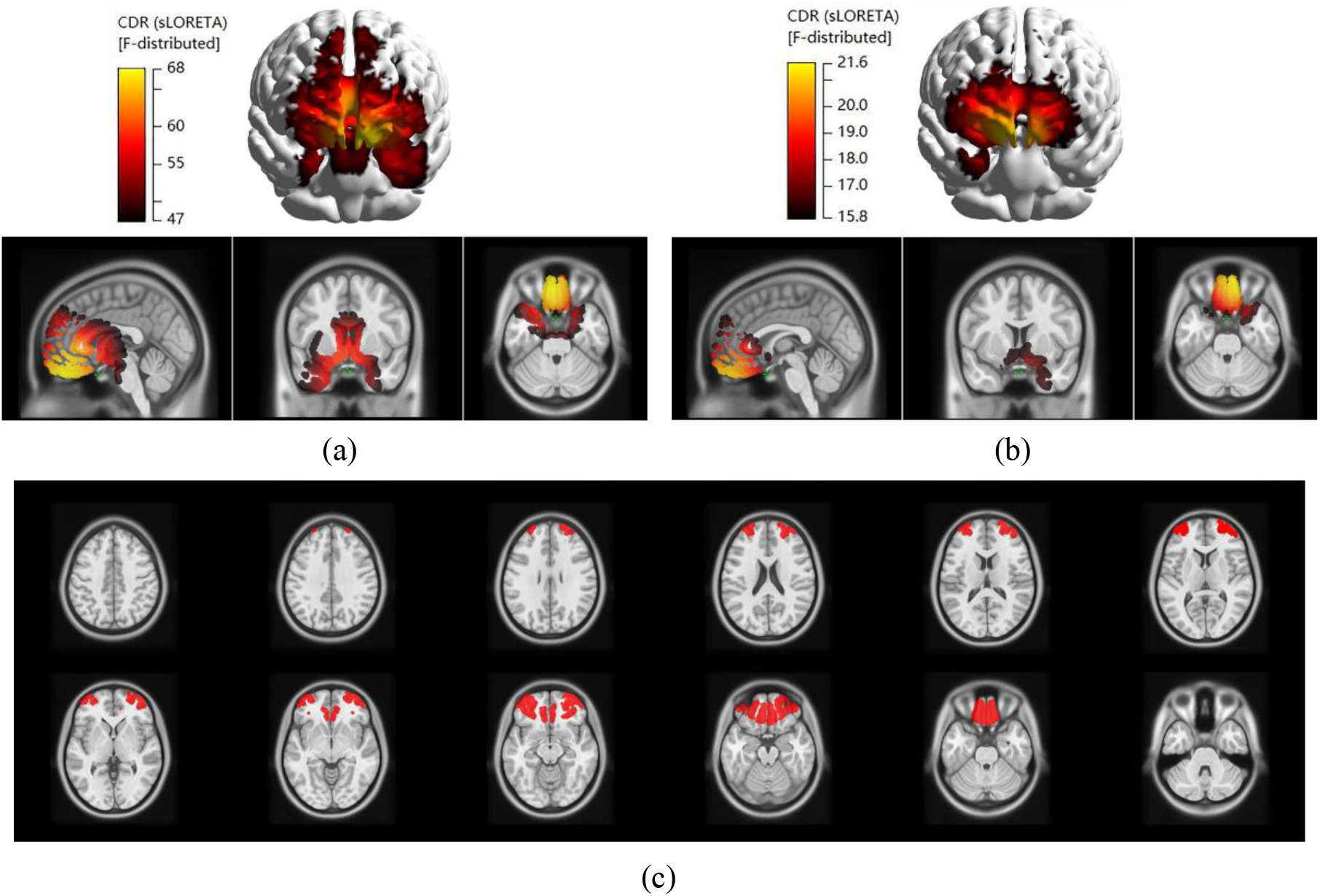
(2a) is the source localization result of the positive group, the maximum current density value F=68; (2b) is the source localization result of the negative group, the maximum current density value F=21.6; (2c) is the differences in source location between positive and negative groups, the red part of the brain slice is the location of the current density difference.

The current density difference of source location is shown in Fig. 2(c). The statistical comparison of source current density is shown in Table 1. The F-value in Table 1 represents the current density value, and the Talairach coordinate is the voxel point coordinate with the largest F-value in voxels. Fig. 2(c) and Table 1 show the difference of source current density between positive group and negative group. The differences can be seen in BA9 (medial frontal gyrus), BA10 (middle frontal gyrus), BA24 (marginal cingulate gyrus),

**Table 1.** Statistical comparison of source current density between positive group and negative group.

| BA area | Brain area | The number<br>of voxels | Maximum<br>F-value of voxel | Talairach coordinate / mm |  |  |
| --- | --- | --- | --- | --- | --- | --- |
|  |  |  |  | *x* | *y* | *z* |
| BA9 | Medial Frontal Gyrus | 57 | 27.72 | -34.44 | 32.17 | 21.79 |
| BA10 | Middle Frontal Gyrus | 27 | 32.43 | -35.36 | 34.77 | 22.11 |
| BA24 | Cingulate Gyrus | 107 | 22.37 | 13.48 | 3.96 | 29.65 |
| BA38 | Superior temporal gyrus | 43 | 22.54 | -50.21 | 1.88 | -6.21 |
| BA45 | Inferior Frontal Gyrus | 29 | 33.63 | -40.17 | 19.95 | 10.35 |
| BA46 | Middle Frontal Gyrus | 84 | 23.76 | 32.13 | 31.4 | 12.75 |

BA38 (superior temporal gyrus), BA45 (inferior frontal gyrus) and BA46 (prefrontal cortex). Among them, BA24 contains the largest number of differential voxels, so the difference of activation range is the most obvious. The difference of voxel F-value in BA45 is the largest, so the difference of activation intensity is the most obvious.

### 3.3. Cortex functional network

#### 3.3.1. Network construction results

The binary correlation matrixes of positive group and negative group are shown in Fig. 3. The abscissa and ordinate in the figure are 64 ROI node numbers. The blue square is 0, indicating no connection. The yellow square is 1, indicating that there is a connection. It can be seen that the positive group has more connecting edges than the negative group.

**Fig. 3.**
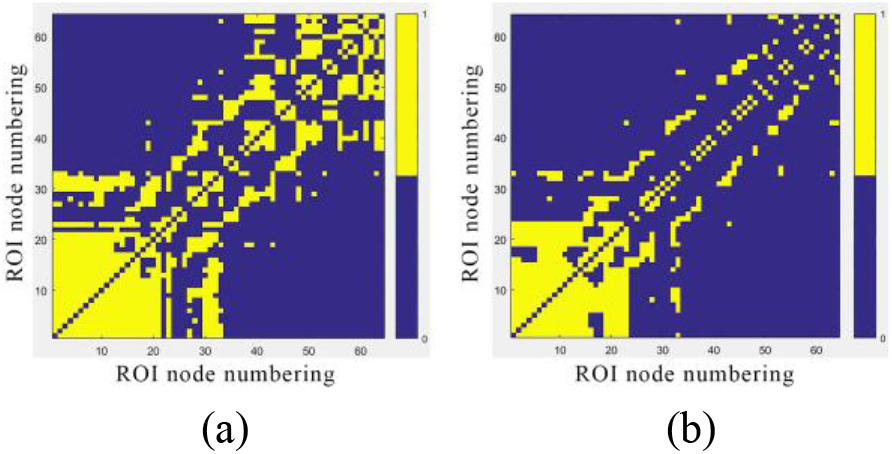
Correlation matrix binary graph of positive group and negative group. (3a) is the binary matrix of positive group; (3b) is the binary matrix of negative group.

Fig. 4 shows the cortex functional network constructed by binary matrix. The red nodes are in the frontal lobe, the orange nodes are in the temporal lobe, the yellow nodes are in the central area, the cyan nodes are in the parietal lobe, the blue nodes are in the occipital lobe, the violet nodes are in the mastoid. If there is a connecting edge between two nodes, it means that there is information transmission between the two nodes. It can be seen that the number of the connecting edges in the positive group is significantly more than that in the negative group. It shows that different emotional states have a significant impact on the functional connectivity of the participants’ brains. Through the comparison, it can be found that the connecting edge of the positive group is inclined to the left hemisphere of the brain, and the connecting edge of the negative group is inclined to the right hemisphere of the brain, but the connecting edges of the brain network in the positive group and the negative group are more concentrated in the frontal lobe.

**Fig. 4.**
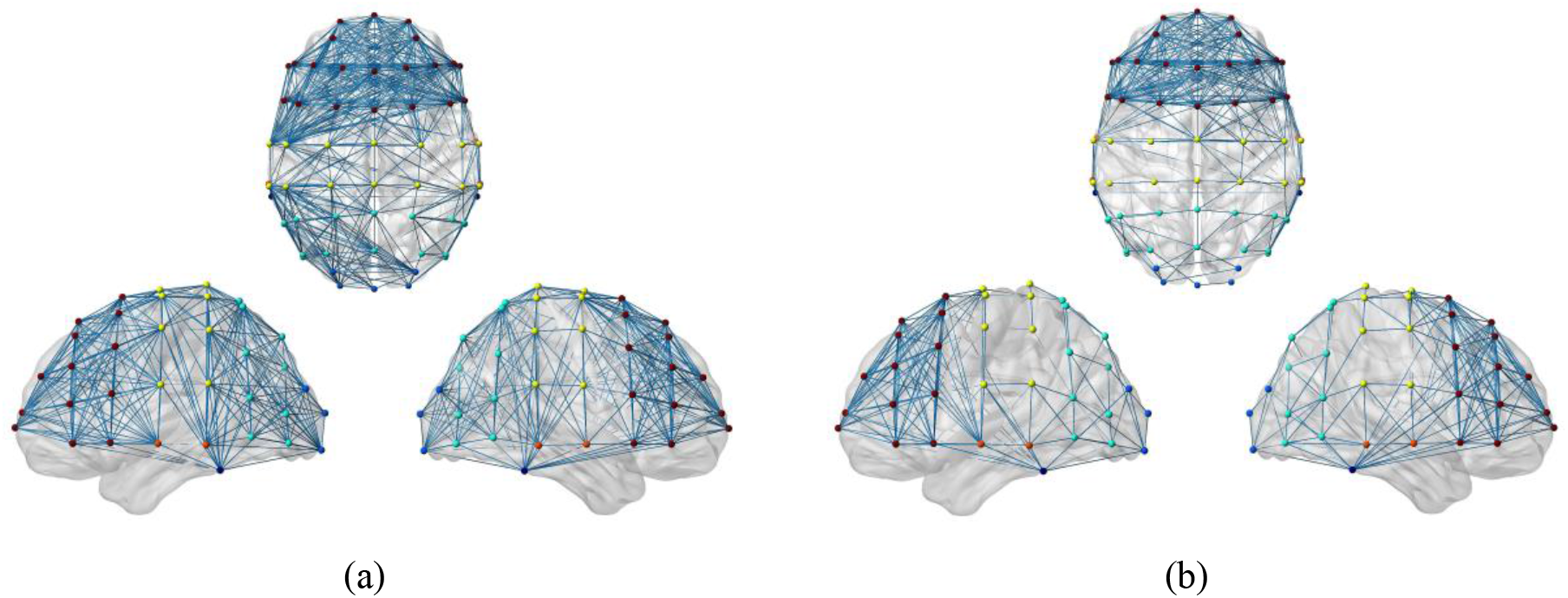
Cortex functional network in positive group and negative group. (4a) is the positive group; (4b) is the negative group. From left to right are left view, top view and right view.

#### 3.3.2. Attribute analysis of the cortex network

In order to further analyze the difference between the positive group and the negative group, we used Pajek 1.0 software to analyze their cortex functional network attributes.

##### 1) Node degree

We used SPSS 20.0 statistical software to conduct two independent sample t-test on the node degree of positive group and negative group, as shown in Table 2. In Table 2, t is the test statistic, sig.(2-tailed) is the significant two tailed *p* value, df is the degree of freedom, Std. Error Difference is the standard deviation of the mean difference of two samples. In addition, the average node degree of positive group is 16.750, and that of negative group is 10.563. The results show that the node degree of positive group is significantly higher than that of negative group (*p*=1.5×10 ^−5^<0.05). It shows that positive emotions stimulate more connections between nodes, which make more information transmitted between nodes, whereas less connections between nodes in negative emotional state.

**Table 2.** Two independent sample t-test results of positive group and negative group.

| Attribute | $t$ | $df$ | Sig.(2-tailed) | Mean Difference | Std. Error Difference | 95% Confidence Interval of the Difference | |
| --- | --- | --- | --- | --- | --- | --- | --- |
|  |  |  |  |  |  | Lower | Upper |
| Node degree | 4.510 | 123.546 | 0.000015 | 6.188 | 1.372 | 3.472 | 8.903 |
| Clustering coefficient | 3.786 | 92.536 | 0.000272 | 0.150 | 0.040 | 0.071 | 0.228 |

##### 2) Clustering coefficient

We conducted two independent sample t-test on the clustering coefficients of positive group and negative group, as shown in Table 2. In addition, the average clustering coefficient of positive group is 0.746, and that of negative group is 0.596. The results show that the clustering coefficient of positive group is significantly higher than that of negative group (*p*=2.72×10 ^−4^<0.05). It shows that the brain network of the positive group is closer and more collectivized than that of the negative group.

##### 3) Global efficiency

We calculated the global efficiency of the cortex functional network in the positive group and negative group. The global efficiencies of positive group and negative group are 0.542 and 0.420, respectively. The global efficiency of the positive group is 28.9% higher than that of the negative group. It shows that the information transmission rate of the positive group is significantly higher than that of the negative group.

##### 4) Small-world property

We calculated the *σ* value of the cortex functional network in the positive and negative groups. The *σ* values of positive group and negative group are 2.294 and 1.765, respectively. It can be seen that the *σ* value of positive group and negative group is greater than 1, which indicate that the brain networks of both groups have small-world property. The *σ* value of the positive group is 29.9% higher than that of the negative group. It shows that the small-world property of positive group is significantly greater than that of negative group.

## 4. Discussion

Through the analysis of behavioral data, we found that under the influence of mixed emotional factors, the incidence of false memory in the positive group was significantly higher than that in the negative group. After the source localization of false memory ERP by sLORETA, we found that the difference of source current density between positive group and negative group was mainly reflected in BA24 (marginal cingulate gyrus) and BA45 (inferior frontal gyrus). According to the neurophysiology researches, BA24 is located in the ventral ganglia of cingulate cortex, which is a part of limbic system. It is connected with amygdala, orbitofrontal cortex and hippocampus, and participates in the emotional system of the brain [29, 30]. BA45, a part of Broca’s area, is located in the frontal cortex and is responsible for semantic tasks and word production [31]. Therefore, we can know that in the identification of keywords, positive emotion prompted participants to stimulate more brain resources, emotional brain areas and semantic brain areas were activated at the same time, which increased the participants’ semantic association of key-words. However, negative emotion inhibited the participants to stimulate brain resources, and the activation location was mainly in emotional brain area, which reduced the participants’ semantic association of keywords. So, the positive group had more false memories than the negative group.

After constructing the cortex functional network for ERP of false memory, we found that the connective edges of positive group and negative group were mainly concentrated in frontal lobe, but the number of connective edges in positive group was more than that in negative group. The frontal lobe is mainly responsible for working memory, information integration and logical reasoning in the brain [32]. The results of network attribute analysis showed that the important information contained in the nodes in the positive group was significantly greater than that in the negative group. The density of brain network, the rate of information transmission and the nature of small world in positive group were also significantly higher than those in negative group. It showed that the participants in the positive emotion state used more mental resources to integrate the information or logical conjecture between the words seen in the learning stage and the words in the deep memory of the brain. Therefore, the positive group produced more false memories than the negative group.

## 5. Conclusions

In this study, memory materials with mixed emotions were used to design false memory experiments. The influence of participants’ emotional state on false memory was studied from three aspects: false memory rate, EEG source localization and cortex functional network. The results show that the incidence of false memory in positive group is significantly higher than that in negative group. In positive emotion state, the frontal lobe of the brain is more active, the connection between the regions in the brain network is closer, and the information transmission speed is faster, which leads to more mental resources for semantic understanding and association of words, and then cause more false memory. However, the frontal lobe of the brain in the negative emotional state are not active, and the brain regions are less connected, which hinders the brain from understanding and associating words, only recalls some of the words they have seen in the learning stage. This paper studies how the emotional state of participants affects the incidence of false memory, which provides a theoretical basis for exploring the working mechanism of the brain. It may be useful for how to better avoid the occurrence of false memory, which has academic value and research significance.

## Ethics approval and consent to participate

The experimental data in this study were obtained with the informed consent of all participants. The institutional review board of the Hebei University of Technology approved the experiment, code HEBUThMEC2020006.

## References

[1] G. Murphy, E.F. Loftus, R.H. Grady, L.J. Levine, C.M. Greene, False memories for fake news during Ireland’s abortion referendum, Psychological Science. 30 (2019) 1449–1459.

[2] Manuel, Anglada-Tort, Thomas, Baker, Daniel, Müllensiefen, False memories in music listening: exploring the misinformation effect and individual difference factors in auditory memory, Memory. 27 (2019) 612–627.

[3] C.J. Brainerd, L.M. Stein, R.A. Silveira, G. Rohenkohl, V.F. Reyna, How does negative emotion cause false memories?, Psychol. 19 (2008) 919–925.

[4] C.J. Brainerd, R.E. Holliday, V.F. Reyna, Y. Yang, M.P. Toglia, Developmental reversals in false memory: Effects of emotional valence and arousal, Journal of Experimental Child Psychology. 107 (2010) 137–154.

[5] J. Storbeck, G.L. Clore, Affect Influences False Memories at Encoding: Evidence from Recognition Data, Emotion. 11 (2011) 981–989.

[6] L. Emery, T.M. Hess, T. Elliot, The illusion of the positive: the impact of natural and induced mood on older adults’ false recall, Neuropsychology Development & Cognition. 19 (2012) 677–698.

[7] W. Zhang, J. Gross, H. Hayne, The effect of mood on false memory for emotional DRM word lists, Cognition & Emotion. 31 (2017) 526–537.

[8] Z. Zheng, M. Lang, W. Wang, F. Xiao, J. Li, Electrophysiological evidence for the effects of emotional content on false recognition memory, Cognition: International Journal of Cognitive Psychology. 179 (2018) 298–310.

[9] M.S. Beato, A. Boldini, S. Cadavid, False memory and level of processing effect: an event-related potential study, Neuroreport. 23 (2012) 804–808.

[10] L. Ding, Y. Lai, B. He, Low resolution brain electromagnetic tomography in a realistic geometry head model: a simulation study, Physics in Medicine & Biology. 50 (2005) 45–56.

[11] R.D. Pascual-Marqui, Standardized low-resolution brain electromagnetic tomography (sLORETA): technical details, Methods Find Exp Clin Pharmacol. 24 (2002) 5–12.

[12] K. Sekihara, M. Sahani, S.S. Nagarajan, Localization bias and spatial resolution of adaptive and non-adaptive spatial filters for MEG source reconstruction, Neuroimage. 25 (2005) 1056–1067.

[13] Y.Y. Kim, Y.S. Jung, Reduced frontal activity during response inhibition in individuals with psychopathic traits: An sLORETA study, Biological Psychology. 97 (2014) 49–59.

[14] M.S. Kim, K.M. Jang, H. Che, D.W. Kim, C.H. Im, 2012. Electrophysiological correlates of object-repetition effects: sLORETA imaging with 64-channel EEG and individual MRI, Bmc Neuroscience. 13, 124. https://doi.org/10.1186/1471-2202-13-124.

[15] H. Aghajani, E. Zahedi, M. Jalili, A. Keikhosravi, Diagnosis of Early Alzheimer’s Disease Based on EEG Source Localization and a Standardized Realistic Head Model, IEEE J Biomed Health Inform. 17 (2013) 1039–1045.

[16] C. Robert, M.R. Iman, R.L. Cannon, 2014. Using quantitative and analytic EEG methods in the understanding of connectivity in autism spectrum disorders: a theory of mixed over- and under-connectivity, Frontiers in Human Neuroscience. 8, 45. https://doi.org/10.3389/fnhum.2014.00045.

[17] Y.H. Jun, T.H. Eom, Y.H. Kim, S.Y. Chung, I.G. Lee, J.M. Kim, Source localization of epileptiform discharges in childhood absence epilepsy using a distributed source model: a standardized, low-resolution, brain electromagnetic tomography (sLORETA) study, Neurological Sciences. 40 (2019) 993–1000.

[18] H. Lv, Y Liu, H. Wang, C. Liu, Z. Wang, Effects of sound therapy on resting‐state functional brain networks in patients with tinnitus: A graph‐theoretical‐based study, Journal of Magnetic Resonance Imaging. 50 (2019) 1731–1741.

[19] J. Wang, Y. Wang, H. Huang, Y. Jia, R. Huang, Abnormal intrinsic brain functional network dynamics in unmedicated depressed bipolar II disorder, Journal of Affective Disorders. 253 (2019) 402–409.

[20] R.A. Clark, N. Nikolova, W.J. Mcgeown, M. Macdonald, 2020. Eigenvector alignment: Assessing functional network changes in amnestic mild cognitive impairment and Alzheimer’s disease, PLoS ONE. 15, e0231294. https://doi.org/10.1371/journal.pone.0231294.

[21] L.J. Jie, S. Marjolein, K. Kaustubh, R. Grega, A. Alan, W.C. Michael, Mapping the human brain’s cortical-subcortical functional network organization, NeuroImage. 185 (2019) 35–57.

[22] Y.Y. Dai, N. Yin, H. Yu, G.Z. Xu, Cerebral cortex functional networks of magnetic stimulation at acupoints along the pericardium meridian, Journal of Integrative Neuroscience. 18 (2019) 79–85.

[23] P.R. Zoladz, D.M. Peters, A.E. Kalchik, M.M. Hoffman, J.N. Talbot, Brief, pre-learning stress reduces false memory production and enhances true memory selectively in females, Physiology & Behavior. 128 (2014) 270–276.

[24] A.D. Patrick, M.M.V. Hulle, fMRI bold signal analysis using a novel nonparametric statistical method, Journal of Magnetic Resonance. 185 (2007) 138–151.

[25] M. Rubinov, O. Sporns, Complex network measures of brain connectivity: uses and interpretations, Neuroimage. 52 (2010) 1059–1069.

[26] J. Lehrer, Neuroscience: Making connections, Nature. 457 (2009) 524–527.

[27] S. Achard, E. Bullmore, Efficiency and cost of economical brain func-tional networks, PLoS Comput Biol. 3 (2007) 174–183.

[28] D.J. Watts, S.H. Strogatz, Collective dynamics of ‘small-world’ net-works. Nature, 393 (1998) 440–442.

[29] G. Miller, The good, the bad, and the anterior cingulate, Science. 295 (2002) 2193–2194.

[30] N.I. Eisenberger, M.D. Lieberman, K.D. Williams, Does rejection hurt? An fMRI study of social exclusion, Science. 302 (2003) 290–292.

[31] C.Y. Wu, E. Zaccarella, A.D. Friederici, Universal neural basis of structure building evidenced by network modulations emerging from Broca’s area: The case of Chinese, Hum Brain Mapp. 40 (2019) 1705–1717.

[32] A. Henri-Bhargava, D.T. Stuss, M. Freedman, Clinical Assessment of Prefrontal Lobe Functions, Continuum: Lifelong Learning in Neurology. 24 (2018) 704–726.

